# Episodic evolution of coadapted sets of amino acid sites in mitochondrial proteins

**DOI:** 10.1101/2020.03.09.983361

**Authors:** A.D. Neverov, G.G. Fedonin, E.A. Cheremukhin, G.V. Klink, A.V. Popova, G.A. Bazykin

## Abstract

The rate of evolution differs between protein sites and changes with time. However, the link between these two phenomena remains poorly understood. Here, we design a phylogenetic approach for distinguishing pairs of amino acid sites that undergo coordinated evolution, i.e., such that substitutions at one site trigger subsequent substitutions at the other; and also pairs of sites that undergo discordant evolution, so that substitutions at one site impede subsequent substitutions at the other. We distinguish groups of amino acid sites that undergo coordinated evolution and evolve discordantly from other such groups. In mitochondrially encoded proteins of metazoans and fungi, we show that concordantly evolving sites are clustered in protein structures. Moreover, the substitution rates within individual concordant groups themselves change in the course of evolution, and phylogenetic positions of these changes are consistent between proteins, suggesting common selective forces underlying them.

**Author summary:** Evolution in most protein sites is constrained by alleles in other sites, this phenomena is called epistasis. Generally, newly arisen allele, if it would not be lost, increases its fitness with time due to adaptive mutations occurred in other sites in the process called the entrenchment. Thus, we expect that the evolution of sites would be coordinated, such that mutations in sites rapidly follow each other on phylogenetic lineages. Indeed, we have observed such coordinated evolution of sites in five proteins encoded in the mitochondria genomes of metazoan and fungi. Unexpectedly, coordinated evolution is observed only for nearby sites on protein structures, such that each protein could be partitioned into several groups of concordantly evolving sites. Evolution of sites from different groups is discordant, i.e. their mutations repel each other into different phylogenetic lineages or clades. Thus, the proteins encoded in mitochondrial genome consist of the sort of structural blocks like elements of the LEGO kit. Some of them have functional specialization, e.g. some blocks are associated with interfaces between proteins composing respiratory complexes.

## Introduction

### Correlated occurrence of amino acids at different sites

The rate at which individual protein sites accumulate substitutions changes in the course of evolution, which violates the assumptions of evolutionary models and may cause problems for phylogenetic reconstruction. This variability can uniformly affect all substitution types (“heterotachy” [1,2]) or differentiate between them (“heteropecilly” [2]). The substitution rate is the product of the rate at which mutations arise and the rate at which they are fixed [3], and can be affected by changes in either of these processes. The main factor affecting the fixation probability is selection favoring some variants over others. The direction or magnitude of selection at a site can change due to multiple forces including changes in environmentally induced constraints or substitutions at other epistatically interacting genomic sites.

The latter process appears to play a major role [4,5]. One type of evidence for this is the correlations between the occurrence of different amino acids at pairs of sites in multiple alignments (MSAs) of homologous sequences. Such correlations, inferred using direct coupling analysis (DCA) or related methods, are associated with physical proximity, and are sufficiently strong that they can be used to infer protein structures and interprotein contacts [6–10] and to predict fitness effects of substitutions [11–13].

Given the success of these approaches, it is tempting to aggregate cooccurrence data across many sites to get a bird’s eye view of the constraints on the evolution of the entire protein. Granata et. al. [14] have processed a DCA-derived cooccurrence matrix by graph vertex clustering. They thus partitioned protein sites into dense domains spatially separated on the structure, and showed that these domains correspond to rigid dynamics domains of the protein. In a different approach, Neuwald [15] partitioned sequences into hierarchically organized subsets, each characterized by a group of sites (domain) conserved within this subset but distinguishing it from adjacent subsets. Comparison of such domains with the protein structure allows to associate them with biological functions [16]. Domains obtained by this approach are only weakly similar to those inferred by factorization of DCA cooccurrence matrices [17], suggesting that these methods can provide complementary perspectives.

Such approaches can reveal groups of sites that are associated with each other, which may help unravel evolutionary constraints linked with some biological functions. However, how these associations manifest themselves in the course of evolution is not understood well.

### Complexity of mitochondrial evolution

Mitochondrial-encoded proteins are a good model system for coevolution between sites. On the one hand, sequencing data is abundant across all eukaryotes, and structures and functions of proteins and protein complexes are well understood. On the other hand, mitochondrial evolution is a complex process. This complexity has been mainly studied from the viewpoint of phylogenetic reconstruction. Mitochondrial proteins violate the basic assumptions of phylogenetic methods, namely, homogeneity of the processes of amino acid substitutions along lineages, between sites in alignment and between character states within sites [18]. This variation can arise due to differences in mutation [19,20] and/or selection [21,22]. As a result, mitochondrial evolution has motivated development of approaches that relax one or several of these assumptions. Heterogeneity between sites has been addressed by using Gamma [23] or general discrete distributions of substitution rates [24] and CAT-models that use Dirichlet processes to classify sites into categories with different equilibrium frequencies of characters [25]. Heterogeneity between lineages has been modeled by using more complicated models [1,2,19].

Another dimension of the complexity of mitochondrial evolution is the high degree of convergence between unrelated lineages. This convergence can be caused by mutational biases [19,26] or natural selection favoring the same amino acid in different lineages at a site (homoplasy) [22,27]. Selection-induced convergence at a small subset of sites may produce erroneous phylogenetic signal overwhelming that from unaffected sites, especially when this convergence at different sites affects the same lineages, which has been observed in all 13 protein-coding mitochondrial genes between the lineages of snakes and agamid lizards [27]. The strong signal of convergent evolution cannot be fully addressed in the phylogeny reconstruction even with modern methods [18].

### Epistasis in mitochondrial proteins

Several lines of evidence suggest that amino acid sites are involved in tight epistatic interactions both within and between mitochondrial proteins [28]. First, sites of mitochondrial proteins COX1, COX2 and COX3 of the cytochrome oxidase complex (COX) that are involved in contact interfaces with other proteins encoded by mitochondrial or nuclear genomes evolve at systematically different rates, compared to sites not involved in such interfaces [29]. The direction of this difference varies among proteins. Evolution is decelerated, indicating stronger purifying selection, at contact interface sites of COX2 and COX3, compared to exposed noncontact noninterface sites. By contrast, in COX1, selective constraint is stronger at non-interface sites, possibly due to their involvement in formation of heme environment. In all three proteins, sites in contact with other mitochondrial-encoded proteins evolve slower than those in contact with nuclear-encoded proteins [29]. These data suggest that the effect of epistasis on the rate of evolution strongly depends on the identity of the interacting partners.

Second, epistasis has been inferred from compensated pathogenic deviations [30], i.e., cases when a human pathogenic variant has been observed in wildtype in some species. Several such cases have been described for the mitochondrial-encoded proteins of oxidative phosphorylation (OXPHOS). Detailed studies have shown that such variants can be neutralized by substitutions at other sites proximal in the 3D structure or in the same interaction interface [31].

Finally, sites in close proximity in protein structures tend to coevolve. This has been observed in COX1 protein [32] as well as in mitochondria-nuclear interfaces [33–35].

Still, our understanding of interactions between sites and the role of such interactions in the evolution of OXPHOS proteins remains limited. Here, extending our previous work [36,37], we develop a phylogenetic method for inference of protein sites involved either in positive or negative epistatic interactions. Roughly, for each pair of sites, we count the number of cases when a substitution at one of these sites rapidly follows a substitution in the other within the same evolutionary lineage. An excess of such cases is suggestive of positive epistasis, whereby the first substitution increases the fitness gain associated with the second substitution; while their deficit is suggestive of negative epistasis, whereby the first substitution decreases the selective advantage, or increases the deleterious effect, of the second one. To address this formally, we compare this number to that expected if substitutions at each site proceed independently of each other. This is done by calculating the association statistic, which is positive if the number of pairs of consecutive substitutions at these sites is unexpectedly high, and negative, if it is unexpectedly low. We apply this method to evolution of OXPHOS proteins encoded in mitochondrial genomes of metazoans and fungi.

In all proteins, we observe many site pairs with positive and negative phylogenetic associations, with the signal of negative associations being much stronger. Using modularity, a community detection method in networks with negative and positive links, we partition sites in each protein into coevolving groups with a high density of positive links within and negative links between groups. Sites within a group tend to be located densely on the protein structure. We show that groups distinguished on the basis of cooccurrence of individual substitutions also demonstrate concordance of substitution rates; and that changes in these rates occur in concert between all five OXPHOS proteins encoded in mitochondrial genomes, suggesting that these changes have a common cause.

## Results

### Concordantly and discordantly evolving pairs of sites

First, we modified the previously developed phylogenetic approach [36] to detect pairs of concordantly evolving amino acid sites in mitochondrial-encoded proteins (fig. 1). In brief, we reconstructed the phylogenetic positions of all substitutions, and identified pairs of sites such that a substitution at one of them frequently triggered a rapid subsequent substitution in the other, as evidenced by higher than expected values of the epistatic statistic [36]. Such pairs were assumed to be concordantly evolving, under the logic that the first substitution increased the selective benefit of the second one, implying positive epistatic interactions. In each of the five studied proteins, we observed strong positive associations of substitutions for a number of site pairs that was significantly above that expected randomly (tab. 1). Surprisingly, at a number of other site pairs, the observed epistatic statistics were significantly lower than expected, indicating that a substitution at one of these sites was followed by a substitution at the other more rarely than expected randomly, and suggesting negative associations whereby the first substitution makes the second one more deleterious. The negative signal was much stronger than the signal of positive associations, both in terms of lower FDR and larger number of significant site pairs (tab. 2). At most of the negatively associated site pairs, the substitutions at the two considered sites tended to occur in lineages that were not only distinct, but also remote from each other on the phylogeny (fig. 2).

**Table 1.** Numbers of concordantly evolving site pairs under different significance thresholds

|  | **p ≤1e-4** |  |  | **p ≤0.001** |  |  | **p ≤0.01** |  |  | **p ≤0.05** |  |  |
| --- | --- | --- | --- | --- | --- | --- | --- | --- | --- | --- | --- | --- |
| gene | #PP |  | FDR | #PP |  | FDR | #PP |  | FDR | #PP |  | FDR |
| ATP6 | 123 | ** | 0.036 | 250 | ** | 0.104 | 683 | ** | 0.357 | 1727 | * | 0.704 |
| CYTB | 221 | ** | 0.068 | 446 | ** | 0.183 | 1313 | * | 0.571 | 3212 |  | 1.02 |
| COX1 | 762 | ** | 0.03 | 1730 | ** | 0.073 | 5199 | ** | 0.227 | 13121 | ** | 0.45 |
| COX2 | 142 | ** | 0.032 | 238 | ** | 0.103 | 576 | ** | 0.408 | 1378 |  | 0.87 |
| COX3 | 306 | ** | 0.019 | 640 | ** | 0.053 | 1680 | ** | 0.183 | 3949 | ** | 0.393 |

For each gene, the number of predicted site pairs (#PP) and the false discovery rate (FDR) for the corresponding nominal p-values are shown. Asterisks indicate the false Exceedance Rate (FER), i.e., the probability that the number of chance-alone findings corresponding to a particular nominal p-value threshold (#FP) is equal or exceeds the number of findings in real data (#PP) (\*, FER≤0.05; \*\*, FER≤0.01).

**Figure 1.**
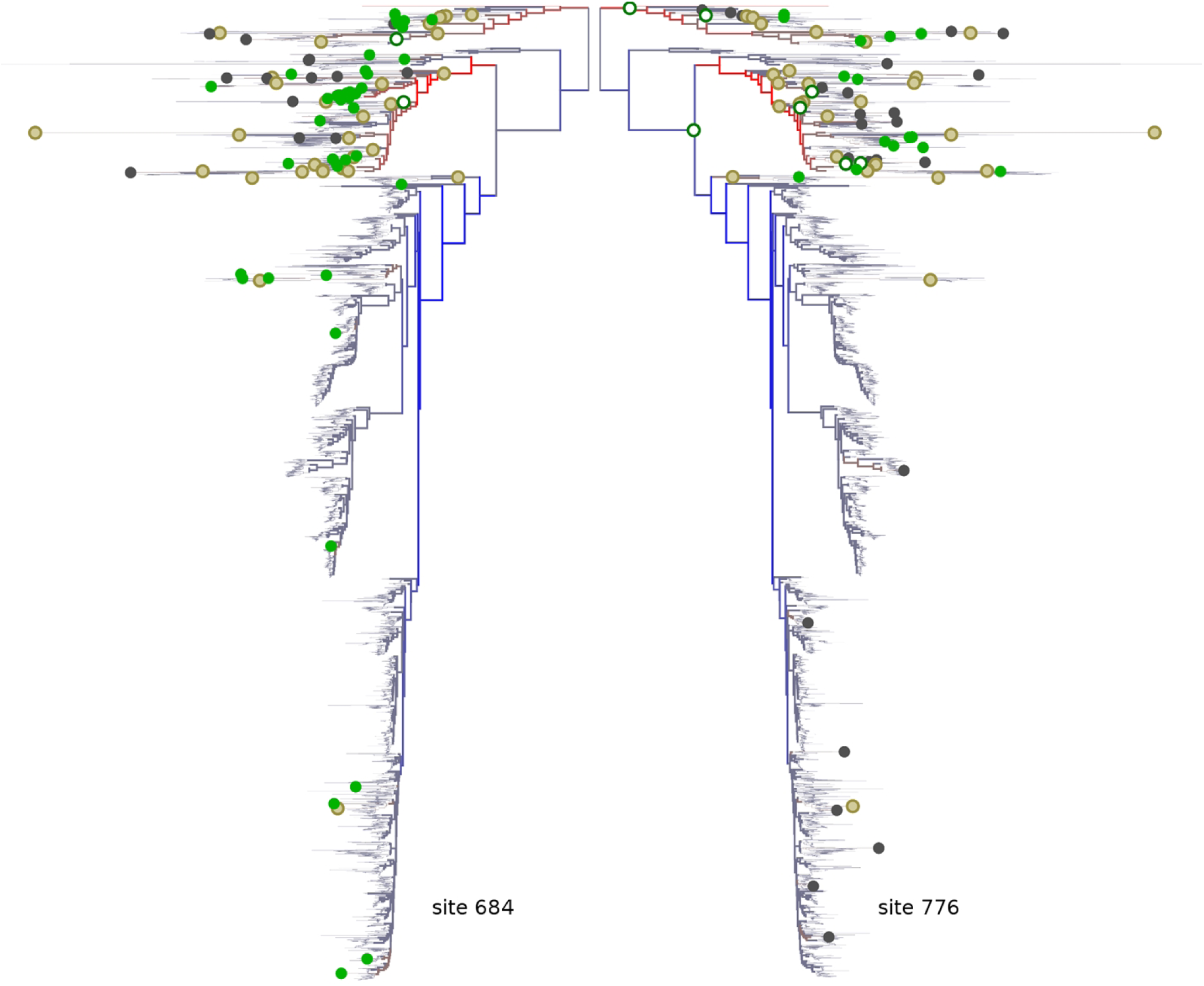
Concordant evolution of protein sites. An example of phylogenetic distribution of substitution densities for a positively coevolving site pair (COX1 protein, sites 684 and 776, both belonging to group 2). For each internal node of the tree, we calculate the number of substitutions in its subtree, and show the differences between the observed and expected numbers of substitutions by colors of branches, with blue indicating a deficit, and red, an excess of substitutions. The expected number of substitutions for a subtree at each site has been estimated as the total subtree length (sum of its branch lengths) multiplied by the corresponding site- or group-specific rate. Individual substitutions are shown with dots: open green dots mark leading substitutions (occurred fist), closed green dots mark trailing substitutions (occurred second), closed sand dots are substitutions at a site that co-occurred with substitutions in another site in the site pair (684,776), and closed gray dots are other substitutions.

**Table 2.** Numbers of discordantly evolving site pairs under different significance thresholds

|  | **p ≤1e-4** |  | **p ≤0.001** |  | **p ≤0.01** |  | **p ≤0.05** |  |
| --- | --- | --- | --- | --- | --- | --- | --- | --- |
| gene | #PP | FDR | #PP | FDR | #PP | FDR | #PP | FDR |
| ATP6 | 1650 | 0.004 | 2453 | 0.01 | 4206 | 0.052 | 6641 | 0.171 |
| CYTB | 2487 | 0.004 | 4061 | 0.013 | 8132 | 0.066 | 14716 | 0.199 |
| COX1 | 1504 | 0.006 | 2635 | 0.02 | 6283 | 0.093 | 13273 | 0.26 |
| COX2 | 764 | 0.004 | 1202 | 0.015 | 2446 | 0.074 | 4522 | 0.218 |
| COX3 | 715 | 0.005 | 1159 | 0.018 | 2385 | 0.091 | 4451 | 0.267 |

For each gene, the number of predicted site pairs (#PP) and the false discovery rate (FDR) for the corresponding nominal p-values are shown. For all genes and nominal p-values, the FER is below 0.01.

**Figure 2.**
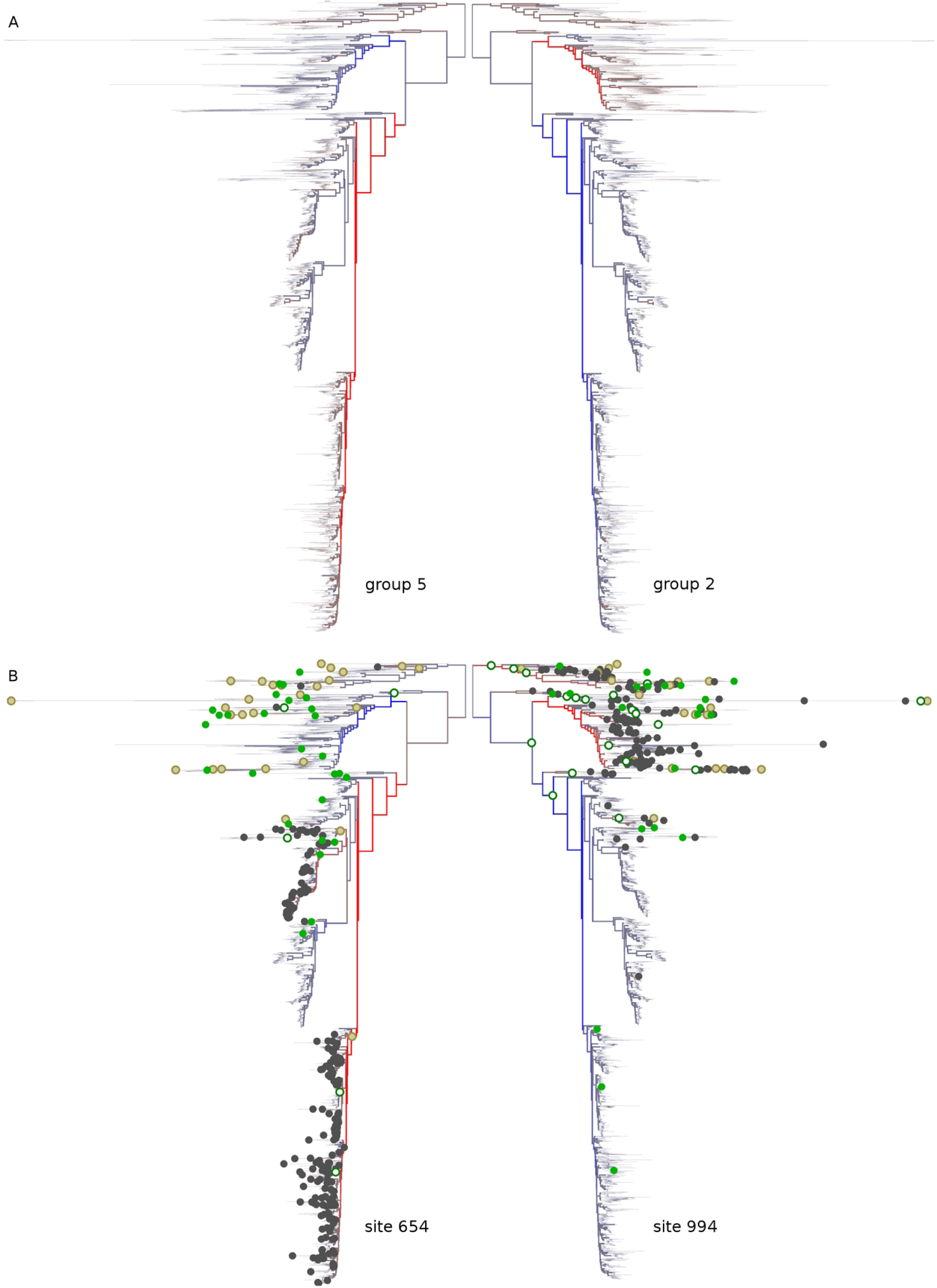
Discordant evolution of protein sites. Examples of distribution of substitution densities for two groups of coevolving sites (a, groups 5 and 2 of the COX1 protein) and for a pair of negatively coevolving sites belonging to these groups (b, site 654 of group 5 and site 994 of group 2). For each internal node of the tree, we calculate the number of substitutions in its subtree, and show the differences between the observed and expected numbers of substitutions by colors of branches, with blue indicating a deficit, and red, an excess of substitutions. The expected number of substitutions for a subtree at each group (a) or site (b) has been estimated as the total branch length multiplied by the corresponding group- or site-specific rate. (a) Groups 5 and 2 (b). In (b), individual substitutions are shown with dots: open green dots mark leading substitutions (occurred fist), closed green dots mark trailing substitutions (occurred second), closed sand dots mark substitutions in a site co-occurred with substitutions in another site in the site pair (654,994) and closed gray dots are other substitutions in each site.

Conceivably, apparent biases in phylogenetic distribution of substitutions, and specifically hemiplasies (spurious convergent or parallel events), could arise from errors in phylogenetic reconstruction. To control for this, we devised a procedure for accounting for uncertainty of phylogenetic reconstruction. Under this procedure, we weight all potentially hemiplasic substitutions corresponding to a single actual substitution by the reciprocal of the number of such hemiplasic substitutions, so that their contribution to the statistic is not inflated (see Methods). Applying this procedure slightly decreased the power of the test: for each nominal p-value threshold, the number of inferred site pairs decreased and the corresponding FDRs slightly increased (tab. S1, tab. S2).

However, this correction changed the list of positively and negatively interacting pairs only slightly; e.g., for COX2, 87 and 72 of the top 100 positive and negative site pairs coincided between the two lists.

### Concordant evolution is associated with proximity in protein structures

While the epistatic statistic provides a useful measure of positive or negative association between substitutions at a pair of sites, its values are not comparable between site pairs. To better understand the patterns of associations between multiple site pairs, for each pair of sites, we converted the epistatic statistic into a normalized form. For this, the statistic for each site pair was z-score transformed, and the resulting z-scores across all site pairs were divided by their maximum value. The resulting values are referred to below as pseudo-correlations; they fall into the range between -1 and 1, with positive values corresponding to coordinated evolution, negative values, to discordant evolution, and 0, to independent evolution of sites.

We asked how the interacting site pairs are positioned relative to each other in 3D protein structures. For significantly positively interacting site pairs (those with positive pseudo-correlations and significant epistatic statistics, see Methods), we found that stronger pseudo-correlations are associated with smaller distances between sites. By contrast, in all genes except COX1, stronger significant negative pseudo-correlations are characteristic of sites remote in the 3D structure; while in COX1, they were characteristic of sites nearby in the 3D structure, similar to positive pseudo-correlations (tab. 3).

**Table 3.** Numbers of concordantly and discordantly evolving site pairs mappable to the protein structures, and correlation between concordance (discordance) and protein structure distance.

| gene | sign of association statistics | #pairs | nominal p-value threshold for FER<0.05 | rho (Spearman's) | p-value |
| --- | --- | --- | --- | --- | --- |
| ATP6 |  |  |  |  |  |
|  | + | 1726 | 0.05 | -0.196 | 2.81E-13 |
|  | - | 6637 | 0.05 | -0.016 | 0.2344 |
| CYTB |  |  |  |  |  |
|  | + | 1511 | 0.0129 | -0.306 | < 2.2e-16 |
|  | - | 14699 | 0.05 | 0.006 | 0.474 |
| COX1 |  |  |  |  |  |
|  | + | 13105 | 0.05 | -0.197 | < 2.2e-16 |
|  | - | 13265 | 0.05 | 0.060 | 2.41E-10 |
| COX2 |  |  |  |  |  |
|  | + | 1009 | 0.0293 | -0.222 | 6.41E-10 |
|  | - | 4513 | 0.05 | -0.053 | 0.002 |
| COX3 |  |  |  |  |  |
|  | + | 3945 | 0.05 | -0.170 | < 2.2e-16 |
|  | - | 4449 | 0.05 | -0.083 | 6.51E-08 |

For each protein, the numbers of significantly concordantly ('+') and discordantly ('-') evolving site pairs with known distances between sites in protein structures (#pairs) are shown, along with the Spearman's correlations (rho) and the corresponding p-value between the association statistic and the structure distance for significant site pairs. For concordantly evolving pairs of sites, a significantly negative value of rho means that strongly associated sites tend to be closer on the structure; for discordantly evolving pairs of sites, a significantly negative rho means that strongly associated sites tend to be apart from each other.

### Detecting groups of coevolving sites

Based on the observed pseudo-correlations, we aimed to construct a coevolution graph in which vertices correspond to individual sites, and edges correspond to either positive or negative associations between them. For this, we transformed the matrix of pseudo-correlations by singling out only significant associations, and, among the positively associated site pairs, only those responsible for direct, rather than spurious, correlations (see Methods). The resulting association statistics were then used to construct the coevolution graphs (tab 4).

**Table 4.**
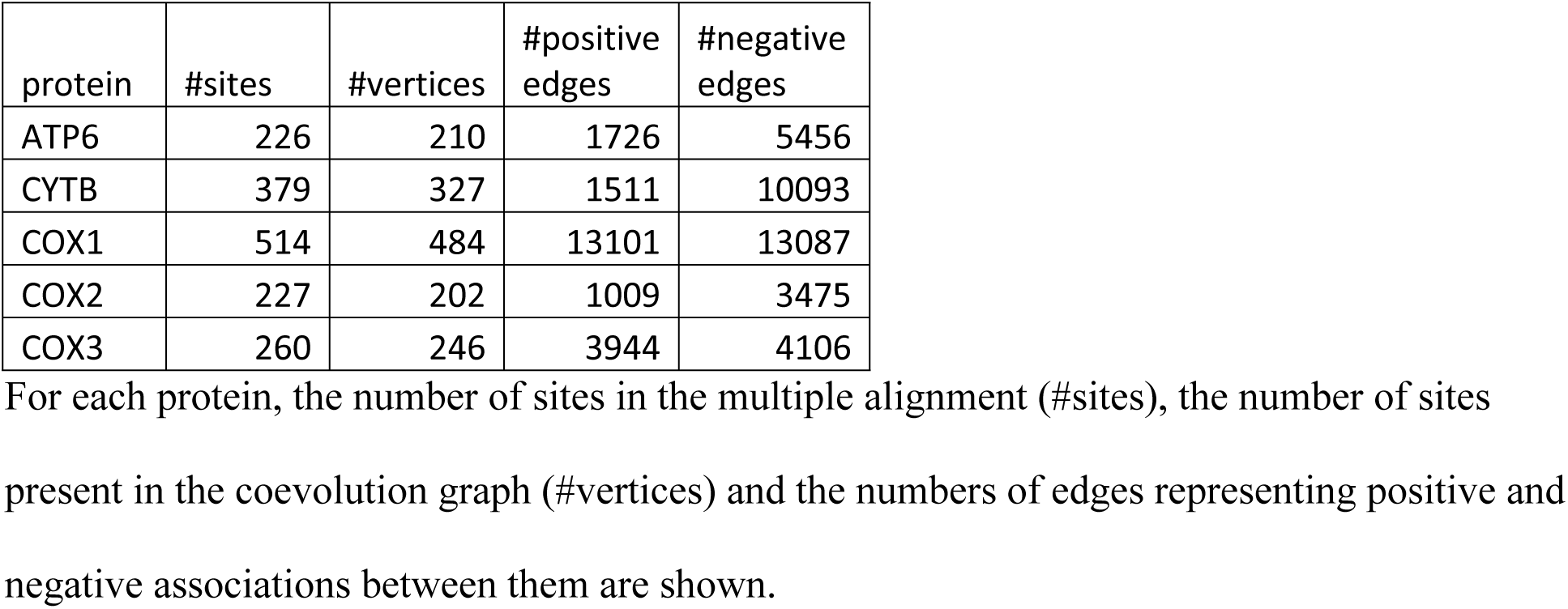
Basic characteristics of coevolution graphs.

The resulting coevolution graphs were then subdivided into subgraphs corresponding to coevolving groups of sites using modularity method for signed graphs [38]. This resulted in groups of sites such that the density of positive edges was high within groups and low between groups, and the density of negative edges was low within groups and high between groups. For each mitochondrial protein, between 4 and 8 groups of sites were thus defined, together including between 82% and 91% of all sites (tab. 5, fig. 3).

**Table 5.** Sites within a coevolving group are colocated on protein structures.

| gene | group | #sites | Observed, in-group contact density | Expected, in-group contact density | p-value |
| --- | --- | --- | --- | --- | --- |
| atp6 |  |  |  |  |  |
|  | 1 | 80 | 0.3 | 0.26 | 0.0471 |
|  | 2 | 71 | 0.29 | 0.23 | 0.0023 |
|  | 3 | 39 | 0.29 | 0.11 | <1e-4 |
|  | 4 | 1 | 0 | 0 | 1 |
|  | total | 191 | 0.45 | 0.35 | <1e-4 |
| cytb |  |  |  |  |  |
|  | 1 | 92 | 0.22 | 0.17 | 0.0009 |
|  | 2 | 62 | 0.14 | 0.11 | 0.0121 |
|  | 3 | 60 | 0.3 | 0.1 | <1e-4 |
|  | 4 | 32 | 0.08 | 0.05 | 0.0378 |
|  | 5 | 29 | 0.16 | 0.05 | <1e-4 |
|  | 6 | 23 | 0.3 | 0.04 | <1e-4 |
|  | 7 | 12 | 0.14 | 0.02 | <1e-4 |
|  | 8 | 6 | 0.04 | 0.01 | 0.0311 |
|  | total | 316 | 0.33 | 0.18 | <1e-4 |
| cox1 |  |  |  |  |  |
|  | 1 | 121 | 0.2 | 0.15 | <1e-4 |
|  | 2 | 112 | 0.23 | 0.13 | <1e-4 |
|  | 3 | 89 | 0.4 | 0.1 | <1e-4 |
|  | 4 | 66 | 0.24 | 0.07 | <1e-4 |
|  | 5 | 39 | 0.08 | 0.04 | 0.0046 |
|  | 6 | 43 | 0.25 | 0.05 | <1e-4 |
|  | total | 470 | 0.39 | 0.19 | <1e-4 |
| cox2 |  |  |  |  |  |
|  | 1 | 49 | 0.18 | 0.15 | 0.0733 |
|  | 2 | 51 | 0.2 | 0.16 | 0.0202 |
|  | 3 | 36 | 0.18 | 0.1 | 0.0009 |
|  | 4 | 36 | 0.24 | 0.1 | <1e-4 |
|  | 5 | 14 | 0.09 | 0.04 | 0.0122 |
|  | total | 186 | 0.32 | 0.22 | <1e-4 |
| cox3 |  |  |  |  |  |
|  | 1 | 67 | 0.2 | 0.16 | 0.012 |
|  | 2 | 63 | 0.21 | 0.15 | 0.0021 |
|  | 3 | 44 | 0.11 | 0.1 | 0.2566 |
|  | 4 | 38 | 0.22 | 0.09 | <1e-4 |
|  | 5 | 25 | 0.1 | 0.05 | 0.0075 |
|  | total | 237 | 0.3 | 0.22 | <1e-4 |

For each protein, we partitioned the vertices in the coevolution graph into groups of coevolving
sites. Contact graph of a protein represents the physical contacts between sites on protein
structures. For each protein, for each group of coevolving sites as well as for entire graph the
mean fraction of edges connecting a site with other sites in the same group, the corresponding expected contact density on random partitions of sites, and the corresponding p-values are shown.

**Figure 3.**
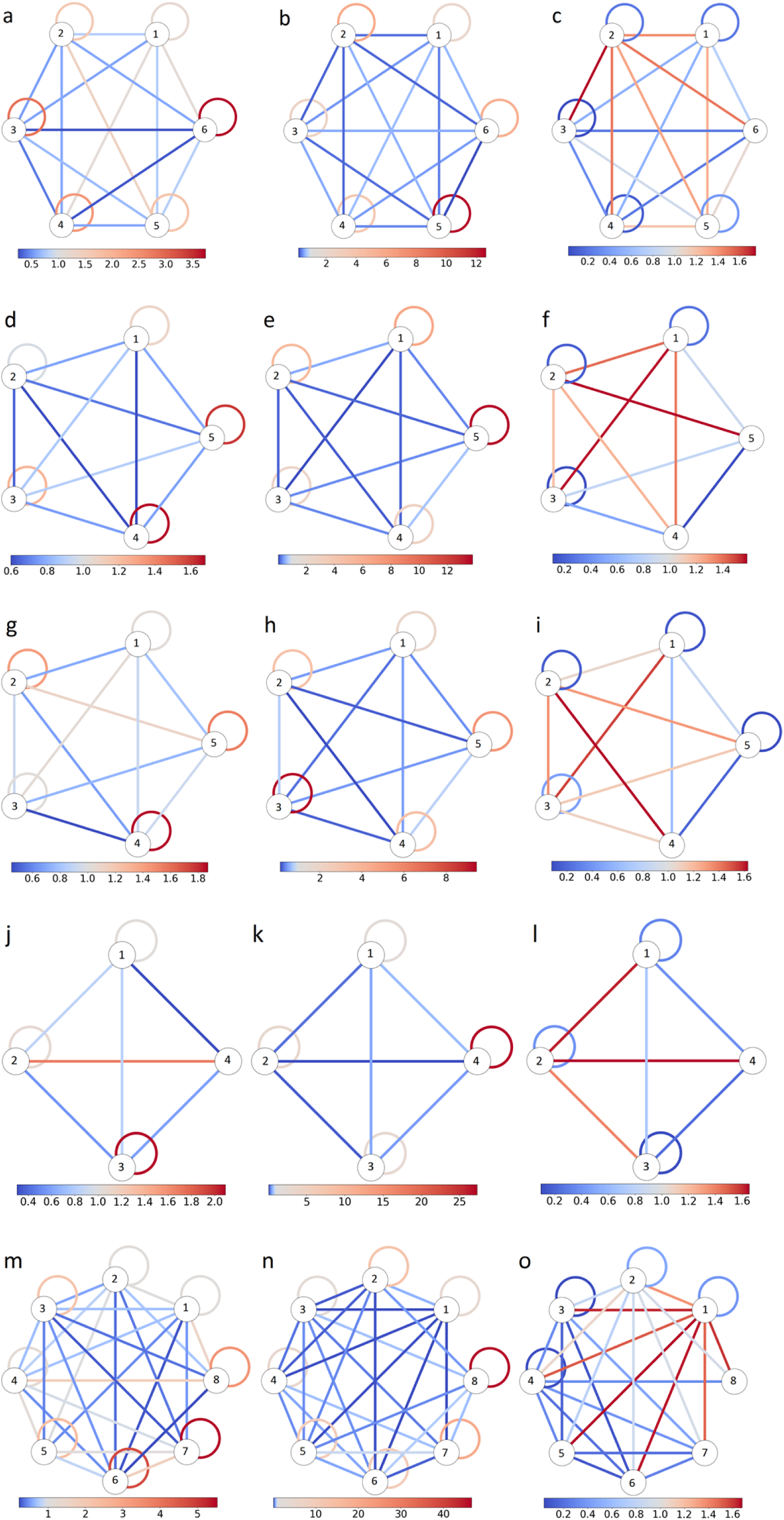
Schematic representation of coevolution and contact graphs for COX1 (a-c), COX2 (d-f), COX3 (g-i), ATP6 (j-l) and CYTB (m-o) proteins. Left column (a, d, g, j and m), contact graphs; middle column (b, e, h, k and n), positive edges in coevolution graphs; right column (c, f, i, l and o), negative edges in coevolution graphs. Each group of coevolving sites is represented by a circle. Links connecting circles represent between-group edges, and links connecting a circle to itself represent within-group edges. The color represent the number of edges (left column) or sum of the weights of edges of the corresponding type (center and left columns) normalized by their expected values obtained from random model used by the vertex clustering algorithm[38]. For contact graphs, groups of coevolving sites are enriched in contacts on the protein structure: the normalized number of edges connecting sites within a group is greater than that between groups. For coevolution graphs, the groups of coevolving sites have larger normalized total weights of positive edges within groups than between groups; by contrast, negative edges tend to have greater normalized total weights between groups.

Next, we asked whether groups of coevolving positions correspond to clusters in the 3D structure of the protein. For this, for each protein, we constructed a second graph, referred to as contact graph. In this graph, vertices again correspond to sites, but there is just one type of edges: two sites are connected if the minimal distance between heavy atoms of their correspondent residues is under 4Å. Considering each group of sites in the coevolution graph of each protein, we then asked whether the corresponding subgraph is tightly connected in the contact graph.

Indeed, coevolving sites were frequently in contact (tab. 5, fig. 3): for each protein, the density of contacts between sites in coevolving groups was higher than expected (P<1e-4). A significantly elevated number of contacts with sites of the same group was also observed for the majority of individual groups of coevolving sites (Table 5), and these groups include the majority of sites (for the p<0.05 threshold, 100% for COX1 and CYTB, 99% for ATP6, 74% for COX2 and 81% for COX3).

### Coevolving groups of sites and interfaces of protein-protein interactions

Next, we tested whether the grouping of sites into coevolving sets tends to be non-random with respect to their involvement in inter-protein interactions with other proteins in the respiratory complexes (either mitochondrial- or nuclear-encoded). In doing so, we controlled for the previously established fact that the coevolving sites tend to be colocalized within a protein. For COX1, COX2,COX3 and ATP6, although not for CYTB, we found that the groups of coevolving sites were non-random with respect to protein-protein interfaces (Fig. 4, 5, Tables S3–S7).

**Figure 4.**
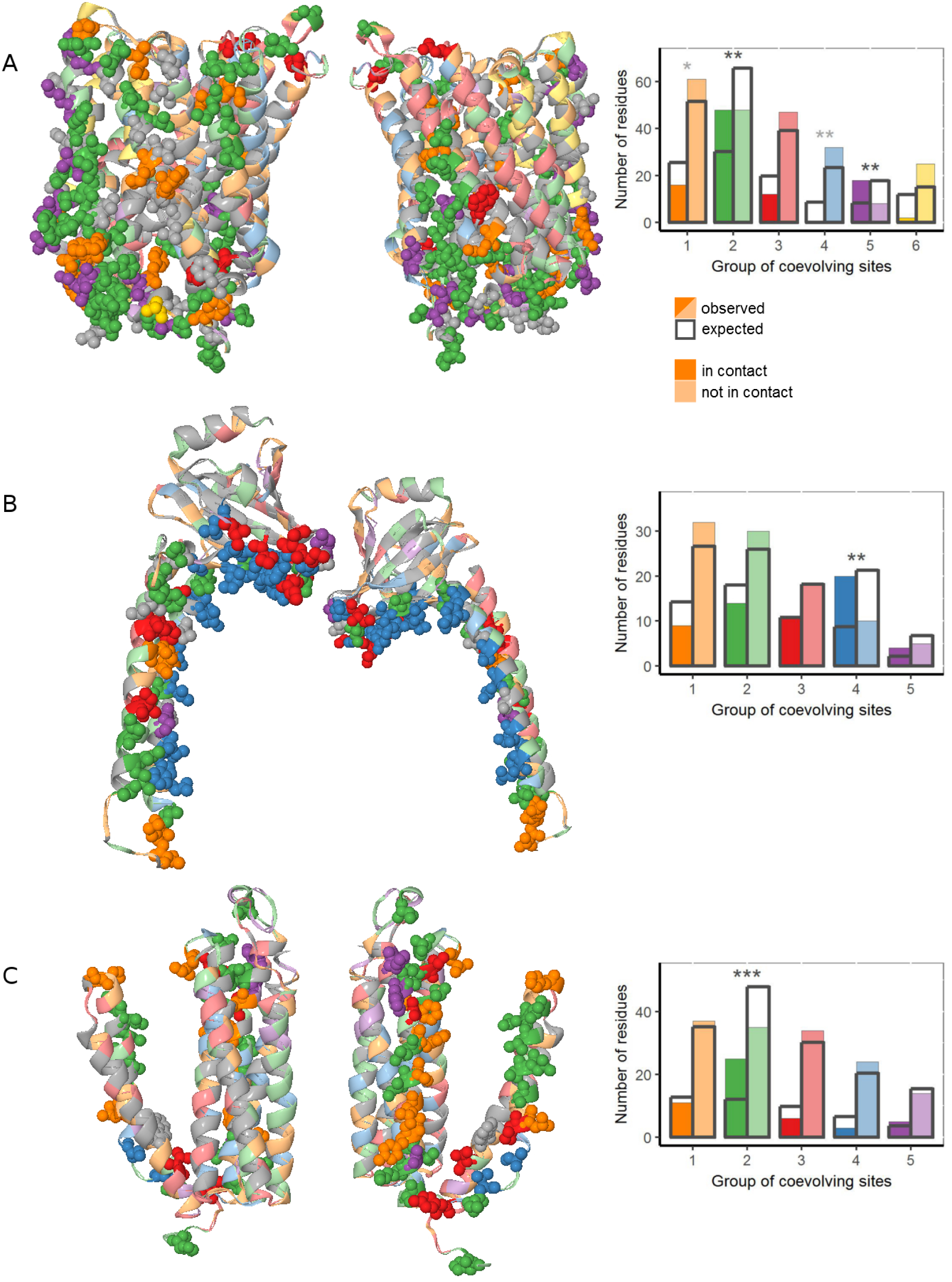
Groups of coevolving sites and interactions between subunits of COX. For each mitochondrial-encoded COX protein: COX1 (a-b), COX2 (c-d) and COX3 (e-f), the groups of coevolving sites are color-coded. (a, c, e) protein structure of COX. Residues at sites involved in interactions with other COX proteins are shown as spheres; residues at sites on protein surfaces not involved in interprotein interactions, as ribbons; the remaining sites of COX were colored in gray. (b, d, f) the numbers of sites which are in contact and numbers of sites which are not in contact with other COX proteins in the protein structure, compared to the expected values. Significant differences are marked with asterisks (*, p<0.025; **, p<0.005; ***, p<0.0005). For COX1 (a-b), interactions with nuclearly encoded COX proteins are considered; for COX2 and COX3 (c-f), interactions with other mitochondrial-encoded COX proteins are considered.

**Figure 5.**
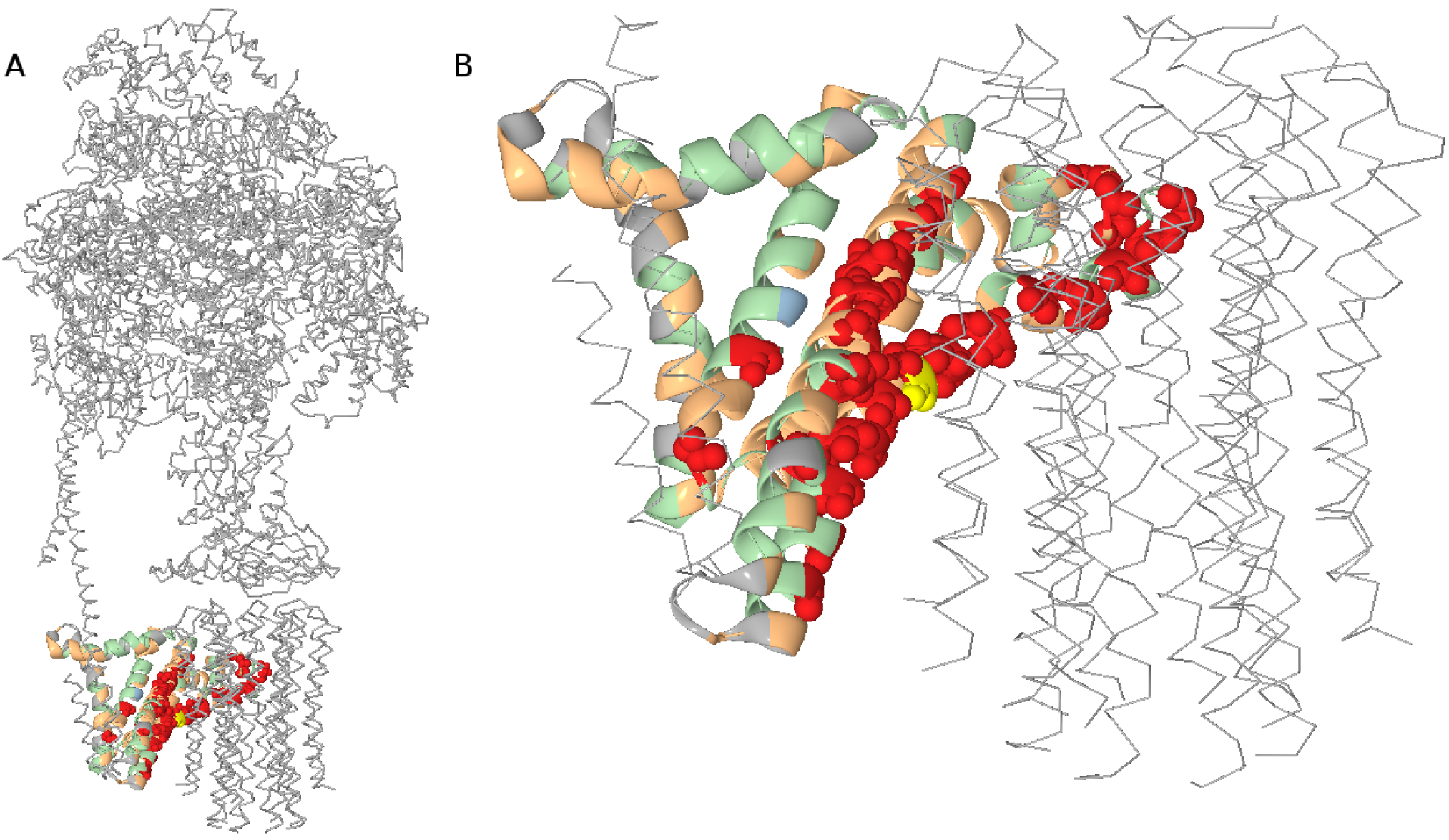
Groups of coevolving sites and function of ATP synthase. For the mitochondrial-encoded ATP6 protein, the groups of coevolving sites are color-coded. Residues at sites of group 3 are shown as spheres; residues at other sites, as ribbons. Most residues at group 3 sites face the rotor part of the ATP synthase and may contribute to proton transport across the membrane. Yellow, Arg159 which also belongs to group 3.

To better understand the link between coevolution and involvement in protein-protein interfaces, we considered each group of concordantly evolving sites in each protein individually. In three of the analyzed proteins, COX1, COX2 and COX3, we found that some groups favored such interfaces, while other groups avoided them (fig. 4, Tables S3–S7). In APT6, most of the sites belonging to the group 3 were located in the helices H5 and H6 and faced the c-ring of the ATP-synthase complex, forming hydrophilic cavities essential for proton transport through the membrane [39]. The conserved arginine 159 crucial for proton translocation [40] also belongs to group 3 (fig. 5).

### Concordant evolution of groups of sites in different OXPHOS proteins

Groups of sites involved in coevolutionary interactions may undergo coordinated acceleration and deceleration of the overall rate of evolution. We aimed to understand when such acceleration or deceleration had taken place. For this, for each protein, we identified a number (between 31 and 94) of branches of the phylogenetic tree out of the total of 4349 internal branches where the relative frequencies of substitutions had changed between groups of coevolving sites, so that the clade of the descendants of this branch has a substitution frequency significantly different from that in the rest of the tree (tab. S8).

We asked whether the identity of such branches was concordant between proteins. To test this, we considered the 2275 branches with enough mutations in coevolving groups to test for a change in mutation frequencies in all five proteins. Since it was impossible to unambiguously position such changes when they had occurred in the two consecutive branches (see Methods), for this test, we shifted the inferred position of each change by one branch towards the root of the tree (i.e., to the parental branch). This resulted in 1611 parental branches that could correspond to frequency shifts at one or both of the daughter branches (tab. 6). Depending on the protein, at between 31 and 90 of these branches, such frequency shifts were actually observed (tab. S8).

**Table 6.** Substitutions rates in groups of coevolving sites have changed concordantly in the evolution of Metazoa and Fungi.

| No. of proteins that have changed their substitution rates concordantly | 5 | 4 or more | 3 or more | 2 or more | 1 or more | 0 or more |
| --- | --- | --- | --- | --- | --- | --- |
| Observed no. of branches with this no. of changes | 5 | 10 | 17 | 45 | 130 | 1611 |
| Expected no. of branches with this no. of changes | 0.1 | 0.8 | 4.2 | 25.1 | 176.8 | 1611 |
| p-value | <1e-4 | <1e-4 | <1e-4 | <1e-4 | 1 | 1 |

We compared the numbers of branches such that the specified number of different proteins (between 0 and 5) changed substitution rates concordantly on this branch. The expected values where obtained from a null-model assuming the same number of rate changes occurring in each protein (tab. S8) independently of the other proteins, with the probability of a change in substitution rates on a particular branch to be proportional to its length.

The identity of the branches corresponding to frequency shifts was unexpectedly similar between proteins (tab. 6). Five branches were concordantly represented for all five proteins; they corresponded to the last common ancestors (LCA) of Fungi and Metazoa, Protostomia and Deuterostomia, Echinodermata+Hemichordata and Chordata, Actinopterigia and Sarcopterigia+Tetrapoda, and Lophotrochozoa and Ecdysozoa. Five branches were each represented for four proteins; these were the LCAs of Cnidaria and Bilateria, Amphibia and Amniota, Otomorpha and Euteleosteomorpha, Neocoleoidea and other taxa within Mollusca, and Neolepidoptera and other taxa within the Holometabola group. 7 branches, including the LCA of Mammalia and Diapsida, were each observed in three proteins; and 28 branches were observed in two proteins.

## Discussion

Epistatic interactions leave a footprint in the evolutionary history of a protein. Explicit reconstruction of past evolution of individual sites allows inferring pairs of sites such that substitutions at them are correlated in time, a pattern which may arise due to positive epistasis. This idea has been used in an approach designed to infer as interacting site pairs those where substitutions occur unexpectedly rapidly after one another [36,37,41,42]. This, however, has only allowed detecting positive epistasis, i.e., the situation in which the first of the two substitutions in a pair increases the selective advantage of the second substitution.

Most existing methods for detection of interactions between sites, such as DCA-based methods, use multiple sequence alignments without explicitly accounting for evolutionary relationships between considered species. Accounting for the phylogeny provides several advantages. First, the MSA-based methods implicitly assume independence of lineages, considering differences in evolutionary distances between lineages as a nuisance factor. By contrast, phylogeny-based methods provide a formal way to account for non-independence between evolving lineages. Second, rooted phylogenies provide explicit polarization of trait states, allowing to distinguish between ancestral and descendant states. In turn, this allows to detect not only positive, but also negative associations between allele pairs.

Here, we make use of this latter advantage. We extend our phylogenetic approach to detect the second possible type of epistatic interactions: negative epistasis. In a pair of negatively epistatically interacting sites, the first substitution reduced the fitness benefit conferred by the second substitution, making the second substitution less probable. As a result, substitutions at negatively epistatically interacting pairs of sites will be “repelled” from one another, leading to a deficit of substitutions occurring one after another at the same lineage.

Any approach based on detection of interactions will have an important caveat which has been discussed previously in the context of positive associations [36,37]. While positively associated substitutions may arise due to interactions, they may also arise due to correlated changes in substitution rates, i.e. due to temporally accelerated mutation or episodes of selection, coinciding between these two sites. Similarly, while a tendency of substitutions to occur at different branches may arise due to negative epistatic interactions, it may also be a result of differences in selection pressure between segments of the phylogeny that differently affect individual sites. Our null model accounts for the differences in selection pressure between branches common to all protein sites, and between protein sites common to all branches. Still, a spurious signal of negative epistasis could arise due to non-epistatically induced changes in selection pressure at a site that are limited to some branches.

We apply our method to the mitochondrial-encoded subunits of the OXPHOS protein complexes. Sites of OXPHOS proteins are known to have undergone changes both in their substitution rates (‘heterotachy’) and spectra (‘heteropecilly’) between lineages. Here, we show that much of this change is correlated between sites, either positively or negatively. Positively interacting sites are positioned close to each other in the protein structure, providing an independent validation for our approach.

Furthermore, by applying community detection methods, we show that the positively interacting sites are grouped into domains of coevolving sites, with negative epistatic interactions distinguishing these groups from each other. Substitutions at sites within the same domain tend to occur in rapid “bursts” on the phylogeny (fig. 1); by contrast, substitutions at sites belonging to different domains tend to occur in distinct clades (fig. 2).

The substitution rate is affected by mutation rate and selection. While the mitochondrial substitution rates have changed due to changes in the mutation rates [19,26], it is unlikely that such changes are site-specific. By contrast, changes in substitution rate may occur naturally at individual sites due to changes in site-specific amino acid propensities. The observed within-site heterogeneity may be most simply explained by such changes in selection with time.

We find that the sites that have changed concordantly are those that are functionally and structurally linked. This is consistent with previous findings obtained by analyzing sequence alignments [9,13,43]. Such concordance may be caused by direct pairwise interactions between sites. Even in the absence of direct interactions, groups of coevolving sites may arise [44] due to protein-level selection forces mediated by one-dimensional, or global, epistasis [45,46].

One possible evolutionary constraint shaping the evolution of COX is the maintenance of interactions between proteins within this complex [47]. Indeed, some of the inferred coevolving groups of sites in COX1, COX2 and COX3 are associated with interactions with other COX proteins, either mitochondrial- or nuclear-encoded or both. Earlier, an excess of biochemically radical substitutions has been observed at interfaces between nuclear and mitochondrial-encoded subunits of COX, suggestive of adaptation [48]. Incompatibilities between nuclear and mitochondrial genomes may play a role in speciation, affecting substitution patterns [47,49,50]; in particular, selection for reproductive isolation may cause bursts of substitutions in interfaces between subunits encoded in nuclear and mitochondrial genomes [33,51,52].

The observed coevolution within interface sites may be partially explained by such selective pressures. To understand what causes changes in substitution rates in groups of concordantly evolving sites, we hypothesized that the evolution of different mitochondrial proteins was also concordant in that evolutionary rate changes were coordinated between different proteins, in addition to their coordination within proteins. Consistently with this hypothesis, we find that different proteins that are subunits of the same as well as different complexes of OXPHOS change substitution rates in groups of sites on coincident branches of the phylogeny. This concordance may reflect changes in the selection pressure on the respiratory function in the process of adaptation to certain ecological niches affecting multiple OXPHOS proteins simultaneously [53]. The branches that had experienced such concordant changes tended to be deeply rooted in the phylogeny, likely indicative of adaptation at the origin of large taxonomic groups. For example, the LCA of mammals indicated concordant changes in substitution spectra for three genes: CYTB, COX3 and ATP6.

Synergistic negative epistasis between deleterious mutations has been described from population genomics data [54] and has been postulated to play a major role in maintenance of sexual reproduction [55]. Furthermore, negative epistasis has been observed in experimental evolution. One characteristic pattern is the presence of distinct “seeding” mutations early in the adaptation process that each trigger its own cascade of subsequent adaptive mutations, directing subsequent evolution. The seeding mutations themselves are in negative epistasis, making them effectively mutually exclusive, and leading to substantial randomness in the choice of the particular adaptive path taken by the population [56]. This pattern is indeed theoretically expected both within and between proteins on fitness landscapes with high local ruggedness [57–59] and has been observed in multiple evolutionary experiments [56,58]. High prevalence of negative epistasis in mitochondrial proteins may have to do with their strong modularity. Conceivably, changes in one domain may trigger subsequent changes in the same domain while increasing the cost of changes in other domains, in line with the “seeding mutations” model [56]. As the respiratory function is carried out by several protein complexes, each consisting of multiple subunits encoded in two genomes with significantly different mutation rates, the negative epistasis between sites or domains of one protein may be driven by the need to support the integrity of this complex system. Our findings that negative epistasis appears to be more prevalent than positive epistasis (Tables 1 and 2), and that positive epistatic interactions tend to be short-range, and negative, long-range in protein structures (Table 3), are also broadly consistent with the results obtained in a high throughput mutagenesis experiment in GB1 protein [60].

Whereas bursts of substitutions in correlated sites caused by positive epistasis have been reported previously [36,37,61,62], negative epistasis has, to our knowledge, not been reported from similar data, probably because detection of a deficit of events is harder than detection of an excess. Families of mitochondrial proteins are an excellent model for the study of negative epistasis between sites because of a high number of substitutions in each site and multiple constraints on their evolution.

## Methods

### Data

Amino acid sequences of five OXPHOS proteins (COX1, COX2, COX3, ATP6 and CYTB) encoded in mitochondrial genomes of 4350 species of metazoans and fungi were obtained from [22]. Each protein was aligned with MAFFT v6.864b [63] using einsi option. For phylogenetic reconstruction, alignment columns with more than 1% of gaps were excluded, and sequences of the five genes were concatenated. The phylogenetic tree was reconstructed with RAxML 8.0.0 using ITOL taxonomy-constrained topology as described in [22]. Bootstrap support for each branch was obtained using rapid bootstrap option of RAxML8.0.0 [64]. For ancestral state reconstruction, we excluded columns with ≥10% of internal gaps. Ancestral states were reconstructed with MEGA-CC using “mtREV with Freqs. (+F)” model and Gamma distributed evolutionary rates between sites with 4 discrete Gamma categories. As the length of COX1 exceeds the limit of MEGA, ancestral states were reconstructed separately for two halves of its alignment.

3D structures were obtained from PDB (1occ for COX, 5ara for ATP6 and 1bgy for CYTB) [65–67]. To map the sites from the MSA to the 3D structure, we performed a pairwise alignment of the *Bos taurus* (TaxID=9913) protein sequence from our MSA to that of the corresponding protein chain in the PDB using BlastP [68,69].

### Inference of epistatic site pairs

To detect epistasis between protein sites, we reimplemented the phylogenetic method from [36], with some modifications, using BioPhylo package for Perl [70]. As in [36], for each pair of sites, we calculated the epistatic statistic as the number of pairs of single amino acid substitutions that were consecutive, i.e., fell onto the same phylogenetic lineage. Mutations that followed one another rapidly had higher weight, with exponential penalties for the waiting time of a second mutation in a pair [36]. Unlike [36], we did not distinguish between “leading” and “trailing” sites; instead, the epistatic statistic was defined for an unordered pair of sites as the sum of the statistics for the two corresponding ordered pairs. As in [36], we compared the observed values of the epistatic statistics with those expected if mutations at different sites were distributed independently of each other, preserving the numbers of mutations for each site and for each branch. To generate these null distributions, we used BiRewire package for R [71]. A total of 10000 sets of mutations were generated in parallel using the GNU Parallel [72] utility. The upper and lower p-values for the epistatic statistic were defined as the percentiles of the null distribution corresponding to the observed values of this statistic.

For each p-value, we estimated the false discovery (FDR) and exceedance (FER) [73,74] rates following the procedure from [36]. Briefly, for 400 random sets of mutations on the phylogeny, we inferred positively and negatively coevolving site pairs. We estimated the FDR as the ratio of the average number of findings (coevolving site pairs with the same or better p-value) in a random dataset to the number of findings in the real data. We estimated the FER as the probability that the number of true positive findings in the data was greater than zero, namely, *P*(#*FP* ≥ #*PP*) < *α*, where #*FP* is the number of false positive findings estimated from a random set, #PP is the number of positive findings in the data and *α* = 0.05 is the significance level. For each gene, we determined the largest sets of significant site pairs as those with positive pseudo-correlations and the nominal p-values of epistatic statistics below the threshold *t*, were *t* is the lesser of the two values: 0.05 and the highest p-value for which FER was <0.05.

To make sure that the observed associations between evolutionary processes at different sites were not artifacts of clustering of spurious substitutions in clades with incorrectly reconstructed topologies [75], we performed a separate analysis accounting for the uncertainty in phylogenetic reconstruction as follows. We defined a subset of well resolved branches of the phylogeny as those with rapid bootstrap [64] support exceeding 95%. These branches split the tree into subtrees with poorly resolved branches. We assumed that the phylogenetic position of the substitutions at well resolved branches was unambiguous. By contrast, the precise number and phylogenetic position of substitutions falling onto a poorly resolved subtree was unknown. We conservatively assumed that each poorly resolved subtree had experienced no more than one substitution at a site. If multiple substitutions within a poorly resolved subtree were reconstructed, we therefore assumed that all but one of these substitutions were spurious. Under this assumption, the phylogenetic position of the only real substitution was not known exactly. We therefore calculated the epistatic statistic as the weighted sum over all of the *n* potential (reconstructed) positions of this substitution within the subtree, each with the weight of 1/*n*.

### Construction of coevolution graphs

For each unordered pair of sites, we defined the pseudo-correlation as the sum of the epistatic statistics for the two corresponding ordered pairs, normalized so that the highest value was 1 if positive, or lowest - 1 if negative. Next, we aimed to single out the site pairs driving the observed positive pseudo-correlations, and to get rid of spurious positive pseudo-correlations resulting from indirect interactions between sites. For this, following previous studies [7,76,77], for each site pair, we defined the association statistic as follows. If the pseudo-correlation was positive, the association statistic was assumed to equal the corresponding partial correlation calculated by cor2pcor R package (http://www.strimmerlab.org/software/corpcor/) with the correlation shrinkage intensity lambda set to 0.9 [78]; if the pseudo-correlation was negative, the association statistic was assumed to equal the pseudo-correlation itself.

For each protein, we then used the values of the association statistic to construct the coevolution graph as follows. All variable sites were represented as graph nodes. We connected a pair of nodes with a “positive” edge if the corresponding site pair had the upper p-value and the corresponding FER both below 0.05 and a positive association statistic. Alternatively, we connected them with a “negative” edge if they had the lower p-value below 0.05 and a negative association statistic. For all five genes, the estimates of FER corresponding to the threshold 0.05 on the nominal lower p-value were below the significance level *α* = 0.05. The values of the association statistic were assigned to each edge as its weight.

To identify groups of coevolving sites, we then applied a vertex clustering algorithm optimizing graph modularity [38] implemented in the louvain package (https://pypi.org/project/louvain/).

### Overlaying coevolution and contact graphs

If the inferred coevolution graphs reflect the structural constraints on proteins evolution, the amino acids adjacent in that graph can be expected to be in spatial contact. To test this, we constructed a contact graph with vertices representing sites, and edges corresponding to contacts in the protein structure. Following earlier studies, we defined a site pair to be in contact if the minimal distance between the heavy atoms of their residues was <4 angstroms [29].

For each group of coevolving sites, we identified the subgraph of the contact graph corresponding to these sites, and defined the contact density statistic as follows. For each group, we calculated the ratio of the number of edges connecting vertices within group to the total number of edges which had at least one vertex in this group. For the entire protein, we calculated the ratio of the number of edges having both vertices in the same group to the total number of edges in the contact graph.

### Associations between groups of coevolving sites and protein-protein interface sites

We estimated the associations between coevolving groups and protein-protein interaction interfaces, defined as follows. Following Aledo et. al. [29,79], we classified the amino acid residues with solvent accessibilities in isolated subunit below 5% as buried; those sites were excluded from the contact graph and not considered further in this analysis. The remaining exposed residues were partitioned into contact residues that had contacts with other subunits in the complex; exposed noncontact interface (ENC_interface) residues that had solvent accessibility within the complex lower than that as an isolated subunit; and the remaining exposed noncontact noninterface residues that were on the protein surface but not involved in interactions with other subunits. Sites in MSA were classified as the corresponding residues of the *Bos taurus* protein. We separately estimated associations of coevolving groups of sites with contact sites and with interface sites, a larger set defined as the union of contact and ENC_interface sites.

The vertex clustering algorithm partitioned protein sites into groups of coevolving sites. We asked whether these groups were enriched or depleted in contact or interface sites, referred to as the testing subsets. To test the null hypothesis of independence, we constructed the contingency table and calculated the chi^2 statistic. Additionally, for each group of coevolving sites, we estimated the Jaccard index, i.e., the number of vertices common to the testing subset and the considered group of coevolving sites, divided by the number of vertices in either of these sets.

Groups of coevolving sites as well as many of the testing subsets formed dense clusters in the protein structure. We were concerned that significant associations between these characteristics could spuriously arise due to such spatial clustering rather than due to interactions between sites. To control for this, we compared the observed values of the statistics with those expected from random groups of sites with the same extent of clustering in the spatial structure as the testing subset. For this, we sampled random subgraphs of the contact graph that had the same number of vertices and equal or greater number of edges connecting them as the testing subset, and used these samples to estimate the expected counts for the contingency tables and to obtain the p-values. To perform this sampling, we implemented an algorithm similar to the algorithm of uniform sampling of connected subgraphs with predefined numbers of vertices [80], with two differences. First, our method rejected subgraphs having fewer edges than the subgraph of the testing subset. Second, we allowed disconnected subgraphs as follows. If the testing subset corresponded to a disconnected subgraph, we performed sampling for each connected component separately, but prohibited the sampled subgraphs corresponding to different components from containing overlapping sets of vertices. For this, we randomly ordered the connected components; sampled a random connected subgraph corresponding to the first component; removed its vertices from the graph; and repeated the procedure for all subsequent components. Sometimes it was impossible to sample a connected subgraph with the given number of edges from the remainder of the graph. To address this, we limited the number of sampling trials to 10000; if no trial succeeded in finding a suitable subgraph, we rolled the algorithm one step back, sampling a different random subgraph for the previous component. Each subgraph thus sampled defined a binary partition of sites.

### Inference of episodic evolution

We analyzed the distribution of substitutions over the phylogeny, identifying the phylogenetic branches corresponding to changes in substitution accumulation rates in some groups of coevolving sites. For this, we used the following procedure. For each branch of the tree *i*, we calculated the vector *n*_*i*_ of the numbers of substitutions that occurred at each group of coevolving sites on this branch, and *m*_*i*_ as the sum of *n*_*k*_ across all branches *k* descendant to *i*. We then traversed the tree, looking for branches corresponding to significant changes in this vector. First, we defined the root branch as the “current ancestor”. Next, starting from it, we traversed the tree towards the terminal branches. For each branch *i* encountered in this process, we compared the vector of substitutions that occurred on this and all subsequent branches *v*_*i*_*=n*_*i*_*+m*_*i*_ to the vector of substitutions that occurred in all the remaining branches descendant to the current ancestor *c*_*i*_=*m*_*a(i)*_-*v*_*i*_, where *a(i)* is the current ancestor to *i*. The two vectors were compared using the Fisher’s exact or chi^2 tests as implemented in the fisher.test package of R, with the Bonferroni correction for the number of internal nodes tested. In each comparison, the groups that had no substitutions in the subtree of the current ancestor were excluded. If *v*_*i*_ and *c*_*i*_ were significantly different, we assumed that the branch *i* corresponded to a significant change in the relative substitution frequencies between groups. In this case, we redefined the current ancestor as *i*, and repeated the procedure for descendant branches. If the total number of substitutions in a subtree of a branch was very low (equal to the number of groups or less), the test was not further applied to descendant branches.

### Estimation of concordance of episodic evolution between OXPHOS proteins

We asked whether the identified phylogenetic branches corresponding to changes in relative substitution frequencies between groups of coevolving sites were coincident between different OXPHOS proteins. For this, first, we obtained the set of branches that were tested for the potential changes in relative substitution rates in all genes, excluding those branches that were not tested in some of the genes because there were not enough substitutions (see above). For each gene, we also identified the subsets of branches corresponding to changes in relative substitution frequencies. In cases when the current ancestor was the immediate ancestor of the tested branches, it was impossible to decide which of the two sister lineages (or both of them) actually experienced the change. Therefore, we positioned the change with the precision of up to two sister branches. Each such pair could be uniquely identified by the name of the parental branch. We tested the significance of the overlap between the branches corresponding to changes in relative substitution frequencies between different genes using a permutation test, assuming that a longer branch was more likely to experience a significant change than a short branch. For this, we calculated the numbers of branches corresponding to changes in zero, one, two, etc., five genes. To calculate the expectation for this value, we generated 10000 permutations, randomly picking for each gene the same number of branches as in the data, each with the probability proportional to the sum of the lengths of its two daughter branches. Finally, we calculated the probabilities to observe the specified number of concordant events for k or more genes, where k=0, 1, 2, …, 5.

## Supplementary Information

Table S1. Numbers of concordantly evolving site pairs, inferred with correction for phylogenetic uncertainty, under different significance thresholds.

Table S2. Numbers of discordantly evolving site pairs, inferred with correction for phylogenetic uncertainty, under different significance thresholds.

Table S3. Coevolution of surface sites of COX1 and interactions with other proteins of the respiratory complex IV.

Table S4. Coevolution of surface sites of COX2 and interactions with other proteins of the respiratory complex IV.

Table S5. Coevolution of surface sites of COX3 and interactions with other proteins of the respiratory complex IV.

Table S6. Coevolution of surface sites of CYTB and interactions with other proteins of the respiratory complex III.

Table S7. Coevolution of surface sites of ATP6 and interactions with other proteins of the respiratory complex V.

Table S8. Substitutions rates in groups of coevolving sites have changed during evolution of Metazoa and Fungi.

The tree, alignments, predicted pairs of epistatically interacting sites and all other data required for analyses mentioned in the Methods section with the detailed description of file contents are provided in the “08_SupplData.7z” file.

